# Gene expression reveals the pancreas of Aselli as a critical organ for plasma cell differentiation in the common shrew

**DOI:** 10.1101/2025.05.22.654563

**Authors:** William R. Thomas, Cecilia Baldoni, Tanya M. Lama, Yuanyuan Zeng, Angelique P. Corthals, Dominik von Elverfeldt, John Nieland, Dina K. N. Dechmann, Liliana M. Dávalos

## Abstract

Almost all mammals rely on the thymus and bone marrow to generate and differentiate B- and T cells essential for adaptive immunity. A few members of the family Soricidae, or true shrews, have also evolved the pancreas of Aselli, a kidney-sized organ hypothesized to serve this primary immune role, and whose gene expression profile is unknown. Here we introduce transcriptomes of juvenile *Sorex araneus* pancreas of Aselli, compare them to those of the spleen and chick bursa of Fabricius, an analogous and bird-specific organ, and explore differential expression overlaps with positively selected genes. While differential gene expression analyses revealed overexpression of genes that regulate the differentiation of B cells into long-term plasma cells (e.g., *IRF4, XBP1, PRDM1*) compared to the spleen and more convergent expression with the bursa of Fabricius than expected by chance (including *IRF4*), overlaps with positive selection were as expected and included *PTPRCAP*, which regulates both T and B cell antigen responses and lymph node size. Our results support the specialized role of the pancreas of Aselli in adaptive immunity, and we propose this unique organ evolved at the intersection between extreme metabolic demands and high parasite burdens in tiny yet very active shrews.

## 1. Introduction

Mammals rely on primary immune organs, such as the thymus and bone marrow, to generate and differentiate cells essential for adaptive immunity, including B- and T cells (*1, 2*). Yet, a few members of the family Soricidae, or true shrews, have evolved a unique immune organ known as the pancreas of Aselli (*3*). This mesenteric lymphoid gland has a dense aggregation of lymphocytes, with a significant proportion made up of large plasma cells in varying degrees of maturation (*4, 5*). In the Eurasian common shrew (*Sorex araneus*), the pancreas of Aselli also accounts for approximately 1% of the total body mass, comparable in size to a kidney (*4*). Based on cellular content, size, and location, it is hypothesized to serve as an additional primary immune organ to defend against their remarkably diverse array of parasites, including helminths (nematodes and cestodes), blood parasites (*Bartonella sp*.), and respiratory fungi (*Pneumocystis carinii*) (*4, 6*–*9*). Despite its novelty among mammals and prominence in shrew physiology, the molecular mechanisms underlying the evolution and function of this immune organ remain largely uncharacterized.

Other clades have evolved similar gut associated lymphatic tissue (GALT) that resemble the pancreas of Aselli (Figure 1A), beyond GALT found in many mammals, like Peyer’s patches (*10*). For example, rabbits possess a more specialized GALT, called the sacculus rotundus, which is a distinct sac-like structure at the terminal portion of the ileum (*11*). However, unlike a true primary immune organ (*12*), the sacculus rotundus does not generate immune cells independently. Instead it is seeded by the thymus (*13*), while also playing an essential role in the hindgut fermentation of plant material (*14*). In contrast, the only comparable analogue to the pancreas of Aselli in evolutionary history is the cloacal bursa (*15*), specifically the bursa of Fabricius when found in birds (*16, 17*). The bursa serves as the primary site of B cell differentiation and proliferation (*18*), where gut-derived antigens that travel through the cloaca help shape antibody diversification. Mature B cells from the bursa then emigrate to peripheral immune organs (*19, 20*), such as the spleen and lymph nodes, contributing immediately toward immune responses. The parallels between the bursa of Fabricius and the pancreas of Aselli are striking; both are unique within their respective clades (shrews and non-ratite birds), both house dense lymphoid populations, and both propagate early-life contributions into developing adaptive immunity.

**Figure 1.**
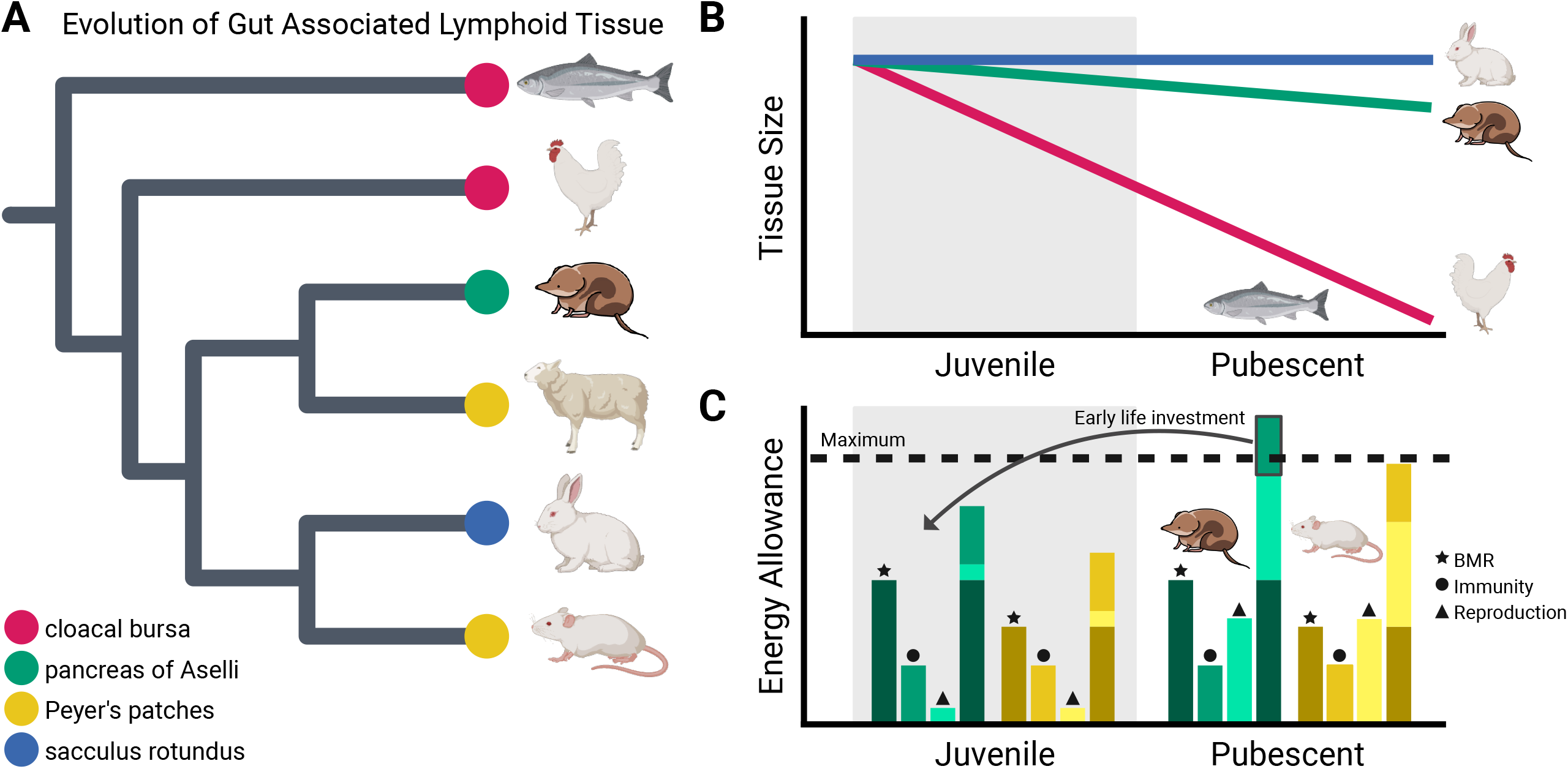
**(A)** Some gut associated lymphoid tissue have independently evolved across vertebrates, giving rise to unique organs such as the pancreas of Aselli in shrews and the sacculus rotundus in lagomorphs. **(B)** Both the pancreas of Aselli and cloacal bursa are prominent early in life, suggesting an important role in early immunity. However, unlike the cloacal bursa, the pancreas of Aselli persists into adulthood, emphasizing function beyond early-life immunity. **(C)** Shrews operate near their maximum energy allowance due to their exceptionally high basal metabolic rates. As they approach reproductive age, their energy demands may exceed sustainable levels, creating trade-offs between physiological functions. The pancreas of Aselli may represent an adaptive strategy that minimizes energetic cost of immune function during reproduction by serving as an early-life investment in immunity.

Despite apparent similarities, development and cellular function indicate that the pancreas of Aselli and the bursa of Fabricius may not be direct evolutionary parallels. One major distinction lies in their developmental trajectories (Figure 1B). The bursa of Fabricius reaches its largest size a few weeks after hatching, but rapidly involutes as birds reach sexual maturity (*16, 21*), much like the thymus in vertebrates (*22*). In contrast, the pancreas of Aselli does not degenerate in size and function upon puberty. Instead, it increases plasma cell storage as the shrew reaches adulthood (*5*). Plasma cells memorize a specific antigen and are longer lived in comparison to mature B cells. Thus, the bursa of Fabricius production of mature B cells is likely for a short-term increase in the number of adaptive immune cells as juveniles, while plasma cell stores in the shrew’s pancreas of Aselli can be used for future antibody secretion as adults.

Life history traits and energetic tradeoffs may help explain why the pancreas of Aselli is unique to shrews and absent in other mammals. Organisms must allocate limited energy between somatic growth, survival, and reproduction, creating evolutionary constraints. Shrews, however, face extreme challenges. With basal metabolic rates nearly twice what is expected for their size (*23, 24*), they have minimal energy reserves and low starvation endurance (*25*), leaving little room for energetic flexibility. Mounting immune responses to their many parasites is especially taxing, particularly during reproduction, which is already energetically demanding for small mammals (*26*). Unlike longer-lived species that can afford to delay reproduction to recover from disease, or short-lived rodents with biannual or continuous breeding shortly after birth, shrews must survive through one winter and then cannot miss their one reproductive opportunity (*27*). Their exceptionally short lifespans compared to most mammals (*28*) restricts them to this single breeding season – meaning that one chance to reproduce is critical. Evolving the pancreas of Aselli may therefore be an adaptive strategy related to reproduction, allowing shrews to invest in long-lived immune cells early in life, before reproduction drains their limited energy reserves, ensuring they can mount efficient immune responses through adulthood (Figure 1C).

We investigated the regulatory functions of the pancreas of Aselli in *Sorex araneus* and its role in plasma cell development by analyzing gene expression. First, we conducted a transcriptomic comparison against a control secondary immune organ, the shrew spleen, analyzing over 20,000 genes to characterize molecular function. Then we compared within-shrew findings to the same analysis conducted with the bursa of Fabricius to assess whether the pancreas of Aselli represents a functional analogue to this avian immune organ. These analyses revealed that adaptative immunity and genes involved in plasma cell differentiation (*XBP1, PRDM1, IRF4*) were significantly upregulated in the pancreas of Aselli compared to the spleen, supporting its specialized role in long-term immune function. While more significant genes with the bursa of Fabricius were found than expected by chance, key genes involved in B cell maintenance (*PAX5, BCL6*) did not overlap, highlighting differences in adaptive immune function. Finally, we explored the effects of selection on the evolution of the pancreas of Aselli by comparing differentially expressed genes against those previously identified as evolving under positive selection in S. *araneus*.

## 2. Methods

### 2.1 Sample collection

*Sorex araneus* were collected (n=4) in November of 2020 from a single German (47.9684N, 8.9761 E) population (protocols authorized by Regierungspräsidium Freiburg, Baden-Württemberg 35-9185.81/G-19/131). Shrews were captured with insulated wooden traps baited with mealworms, which were monitored every two hours to reduce trap-related stress and food insufficiency on gene expression. Once trapped, shrews were aged and sexed (two male and two female) based on tooth wear, fur condition, and gonad development. Notably, Eurasian common shrews have a single breeding season in late spring or early summer and typically do not survive in the wild beyond their second fall. Thus, all shrews captured were prepubescent individuals approximately six months of age. Shrews were euthanized with PAXgene Tissue Fixative through vascular perfusion. Then the pancreas of Aselli and spleen were removed and placed in PAXgene Tissue Stabilizer for 2-24 hours. These organs were then frozen and stored in liquid nitrogen (−180°C).

### 2.2 RNA extraction and sequencing

RNA was extracted from each organ using a modified version of the Micro RNeasy Micro kit (Qiagen). Modifications included 1) tissues were ground using glass mortar and pestles on dry ice for 1-2 minutes to reduce heat-associated RNA degradation, 2) column binding efficiency and lysing was improved by adding 7μL of 2M dithiothreitol and 5μL of 4ng/μL carrier RNA to the lysate, 3) lysate was ground for an additional minute and homogenized with QIAshredder columns, and 4) DNAse incubation was reduced from 15min to 2min. Azenta Life Sciences performed quality control analyses (nanodrop and RNA ScreenTape), library preparation (poly-A selection), and sequencing (approx. 15-25million reads/sample, 150bp PE) of the spleen, while the BGI (Beijing Genomics Institute, California) conducted these analyses for the pancreas of Aselli.

### 2.3 Differential expression

Raw reads for each sequencing run were analyzed with fastp (*29*) to trim adapters and remove low quality sequences. Sequences were then aligned to the *S. araneus* transcriptome (GCF_027595985.1_mSorAra2.pri) using Kallisto 0.46.2 (*30*). Read counts were normalized using the median of ratios from DESeq2 (*31*) which accounts for library content and size. This step was critical to our analysis as organs were sequenced at different sequencing companies. We then ran a principal component analysis (PCA) using the 5,000 genes with the most variation in the data set. Next, we tested for differential expression between the spleen and the pancreas of Aselli samples using DESeq2 (*31*), where gene (n=24,205) counts are fitted with a negative binomial generalized linear model and then tested for differential expression using a Wald test, using sex as an additional covariate (∼sex+organ). The spleen was chosen as a control because, as a secondary immune organ, it does not primarily generate immune cells. This allowed us to identify differential gene expression related to this primary immune role that are specifically upregulated in the pancreas of Aselli. Significant results are identified following a multiple test correction (Benjamini Hochberg; p_adj_<0.05) (*32*). DESeq2 results were then run through a ranked Gene Ontology (GO) enrichment using Kyoto Encyclopedia of Gene and Genomes (KEGG) pathways with the fsgea package (*33*) to explore the molecular functions of differentially expressed genes. We then examined which genes that were upregulated in the pancreas of Aselli overlapped those that evolved under positive selection from previous comparative analyses (cite). A hypergeometric test (n=100,000) was used to test if overlap in significant upregulated genes with the bursa of Fabricius, while a Chi-squared test was used to test if observed overlap with positively selected genes, were greater than expected by chance (p<0.05).

To test if the bursa of Fabricius is a functional analogue of the pancreas of Aselli, we required age-matched transcriptomic data from the bursa of Fabricius and spleen in a bird species. RNA sequencing data for these tissues were obtained from the Sequencing Read Archive (SRA) on NCBI (NIH National Center for Biotechnology Information). We searched the SRA using the keyword “bursa” and restricted results to RNA-seq data from chickens (*Gallus gallus*). Combining three experiments (BioProject PRJNA494531, PRJNA623599, PRJNA511787) provided age-matched control spleen (n=3) and bursa of Fabricius (n=5) samples. Although these chickens were prepubescent, akin to the shrews, they were only two to three weeks old, whereas the shrews were six months old. Older bursa of Fabricius samples exists, however with a sample size that cannot measure variation (PRJNA901530, n=2). This age difference introduced a potential confounding variable in our comparative analyses. To process the sequencing data, we trimmed adapter sequences and removed low-quality reads using fastp (*29*). The cleaned reads were then mapped to the chicken transcriptome (GCF_000002315.5_GRCg6a) using Kallisto 0.46.2 (*30*). Read counts were normalized using the same approach as for *S. araneus*, and differential expression between the spleen and bursa of Fabricius was assessed using DESeq2 (∼breed+organ), with p-values corrected via the Benjamini-Hochberg method (*32*).

## 3. Results

### 3.1 RNA sequencing and alignment

RNA sequencing of *S. araneus* and *G. gallus* organs yielded high-quality data with comparable sequencing depths across organ type and species. *S. araneus* samples had a mean post-filtration sequencing depth of 41 million reads, with sequencing depths within an order of magnitude between the spleen (mean 34.8 million reads) and the pancreas of Aselli (48.1 million reads).

These results were consistent with publicly available transcriptomic datasets on the SRA. For *G. gallus*, the mean sequencing depth was 58.5 million reads, with specific BioProjects showing averages of 53.4 million reads for PRJNA494531 (bursa of Fabricius), 65.6 million reads for PRJNA511787 (bursa of Fabricius), and 58.98 million reads for PRJNA623599 (spleen). Mapping efficiency of sequencing data was similarly high across species. On average, 71.1% of *S. araneus* reads mapped to the NCBI transcriptome, with spleen samples achieving 77.5% mapping, slightly higher than the pancreas of Aselli mapping at 64.7%. *Gallus gallus* mapping rates were comparable to those of the common shrew, averaging 77.3% for the spleen and 79.0% for the bursa of Fabricius.

### 3.2 Differential expression

Thousands of genes were differentially expressed between the *S. araneus* pancreas of Aselli and spleen, with several immune-related pathways – particularly those associated with adaptive immunity – upregulated in the pancreas of Aselli. To assess overall transcriptomic differences between these organs, we first performed a PCA (Figure 2). This analysis revealed a strong separation between the two organ types along PC1, while intra-organ variation, particularly within the spleen, was captured on PC2. Given this clear distinction in gene expression profiles, we proceeded with a differential expression analysis to identify specific genes promoting these differences. Of the 24,205 genes tested, 5,997 were significantly differentially expressed after multiple test correction (p_adj_<0.05), with 2,500 upregulated in the pancreas of Aselli and 3,497 downregulated. Genes critical to plasma cell differentiation were significantly upregulated – *XBP1* (p_adj_<0.001, Log Fold Change=1.67, [LFC]), *PRDM1* (p_adj_<0.001, LFC=1.76), and *IRF4* (p_adj_<0.001, LFC=0.95), while most key genes in B cell maintenance were not – *PAX5* (p_adj_=0.86, LFC=-0.08), *BCL6* (p_adj_=0.31, LFC=0.38), *BACH2* (p_adj_<0.001, LFC=1.70) (Figure 2). To further determine the functional significance of these differences, we conducted a ranked pathway enrichment analysis of KEGG pathways. This analysis identified 22 enriched pathways (p_adj_<0.05), with 11 upregulated and 11 downregulated in the pancreas of Aselli (Figure 2). Notable downregulated pathways included porphyrin and chlorophyll metabolism (p_adj_<0.001, NES=-2.17), while key upregulated pathways were predominantly immune-related. These included primary immunodeficiency (p_adj_<0.001, NES=2.03), antigen processing and presentation (p_adj_<0.001, NES=2.02), T cell receptor signaling (p_adj_<0.001, NES=1.65), natural killer cell mediated cytotoxicity (p_adj_<0.05, NES=1.46), and cytokine-cytokine receptor interaction (p_adj_<0.01, NES=1.45).

**Figure 2.**
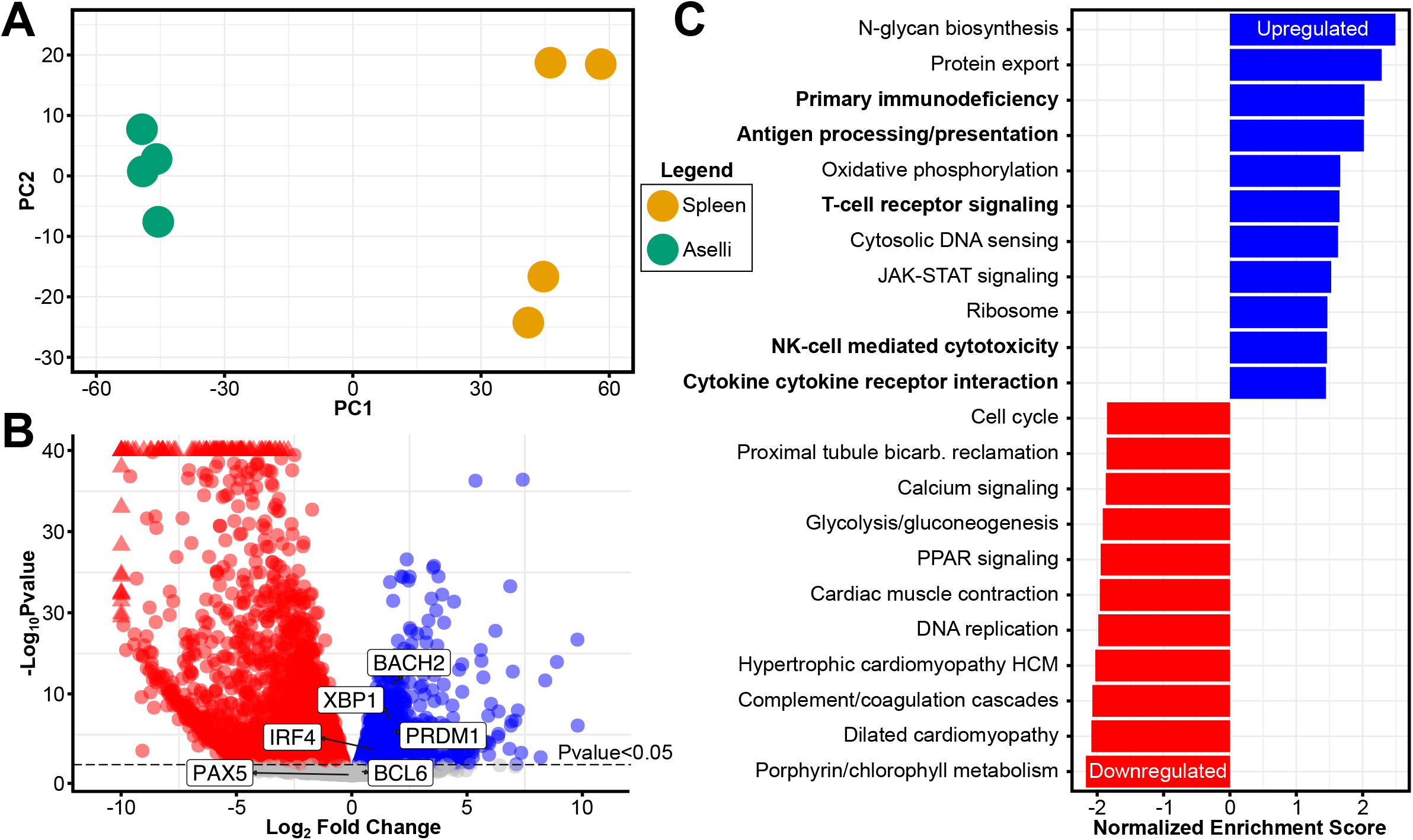
**(A)** Principal component analysis (PCA) of the 5000 most variable genes shows distinct separation between the spleen and pancreas of Aselli along PC1. **(B)** Volcano plot quantifying differentially expressed genes (p_adj_<0.05), with 2,500 significantly upregulated genes (blue) and 3,497 downregulated genes (red) in the pancreas of Aselli. Noted gene include those associated with B cell maintenance (*PAX5, BCL6, BACH2*) and plasma cell differentiation (*XBP1, PRDM1, IRF4*). **(C)** Pathway enrichment analysis revealed 11 upregulated (blue) and 11 downregulated (red) pathways in the pancreas of Aselli, suggesting adaptive immune-related function.

Comparisons to positively selected genes identified overlap with differential expression and selection. Of the 2,500 genes upregulated in the pancreas of Aselli and the 231 inferred to be under positive selection from previous work (*34*), 21 were found to overlap between both data sets. These genes notably include the plasma cell differentiation gene *PRDM1*, as well as the protein tyrosine phosphatase receptor type C-associated protein, *PTPRCAP*. However, this overlap was not greater than expected by chance 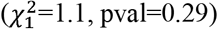.

To test whether the bursa of Fabricius is a functional analogue of the pancreas of Aselli, at least at the transcriptomic level, we performed a differential expression analysis of the chicken bursa of Fabricius to the spleen and compared these results to the same analysis in the common shrew. Of the 23,747 genes tested, 11,184 were significantly differentially expressed, with 5,744 upregulated and 5,440 downregulated in the bursa of Fabricius. Among these, 964 genes were significantly downregulated in both the pancreas of Aselli and bursa of Fabricius and 543 upregulated in both. This overlap is significantly more than expected by chance (hypergeometric test, p<0.001). Next, we examined the genes critical to plasma cell differentiation (*XBP1, PRDM1, IRF4*) and B cell maintenance (*PAX5, BCL6, BACH2*). All three of the B cell maintenance genes were upregulated in the *G. gallus* bursa of Fabricius - *PAX5* (p_adj_<0.001, LFC=3.08), *BCL6* (p_adj_<0.001, LFC=1.01), and *BACH2* (p_adj_<0.001, LFC=1.61). However, only two plasma cell differentiation genes, *XBP1* (p_adj_<0.001, LFC=-0.48), *PRDM1* (p_adj_<0.001, LFC=0.38), *IRF4* (p_adj_<0.001, LFC=1.14), were significantly upregulated, which were all upregulated in the common shrew pancreas of Aselli (Figure 3).

**Figure 3.**
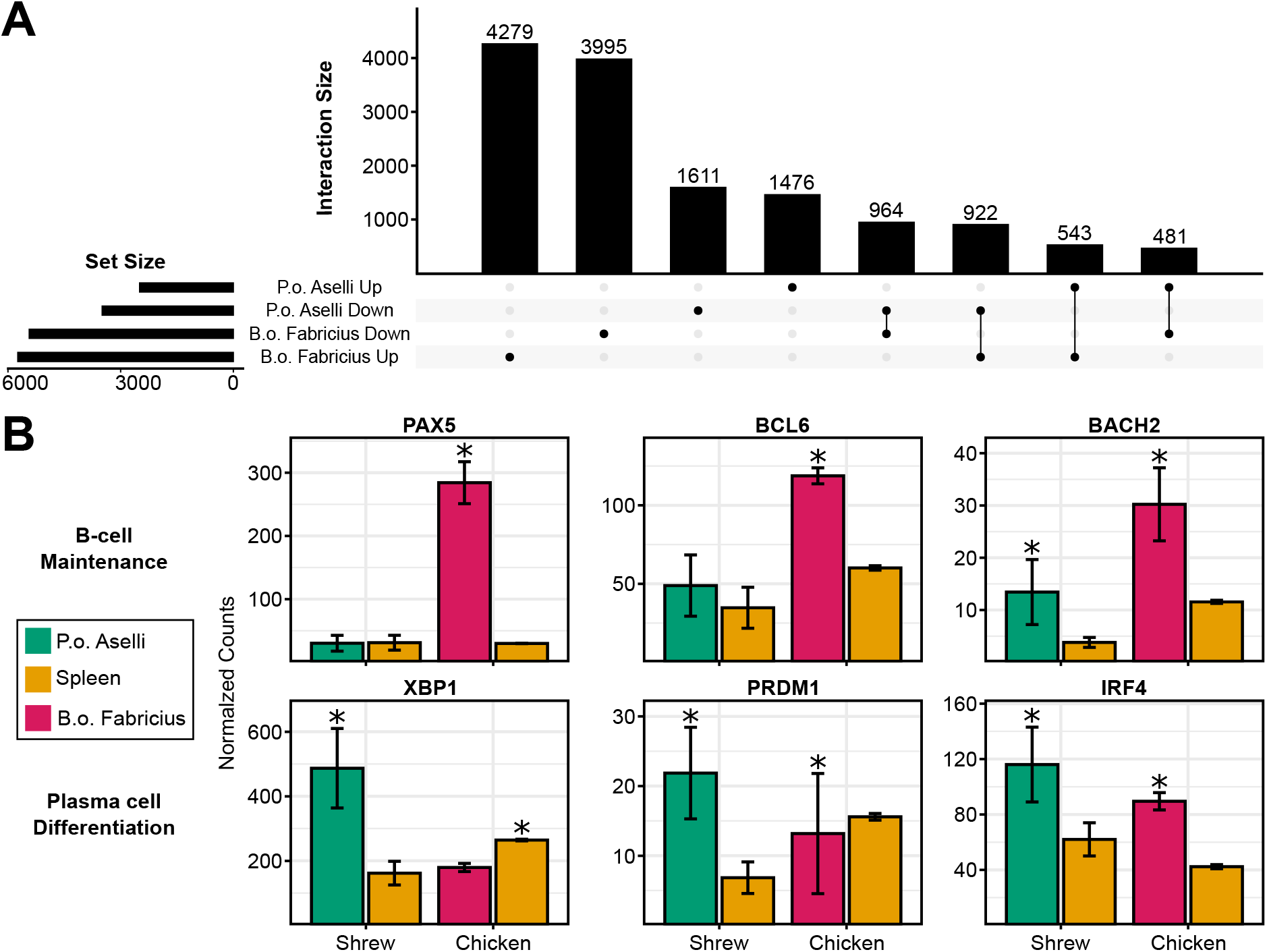
**(A)** Upset plot showing overlap in significantly differentially expressed genes between the pancreas of Aselli vs. spleen in *Sorex araneus* and the bursa of Fabricius vs. spleen in *Gallus gallus*, with 1,507 genes exhibiting concordant regulation. **(B)** Comparative analysis of important B cell differentiation genes (*PAX5, BACH2, BCL6*) and plasma cell differentiation genes (*XBP1, PRDM1, IRF4*) reveals differences in regulatory patterns between the shrew’s pancreas of Aselli and the chicken’s bursa of Fabricius, suggesting divergent adaptive immune function.

**Figure 4.**
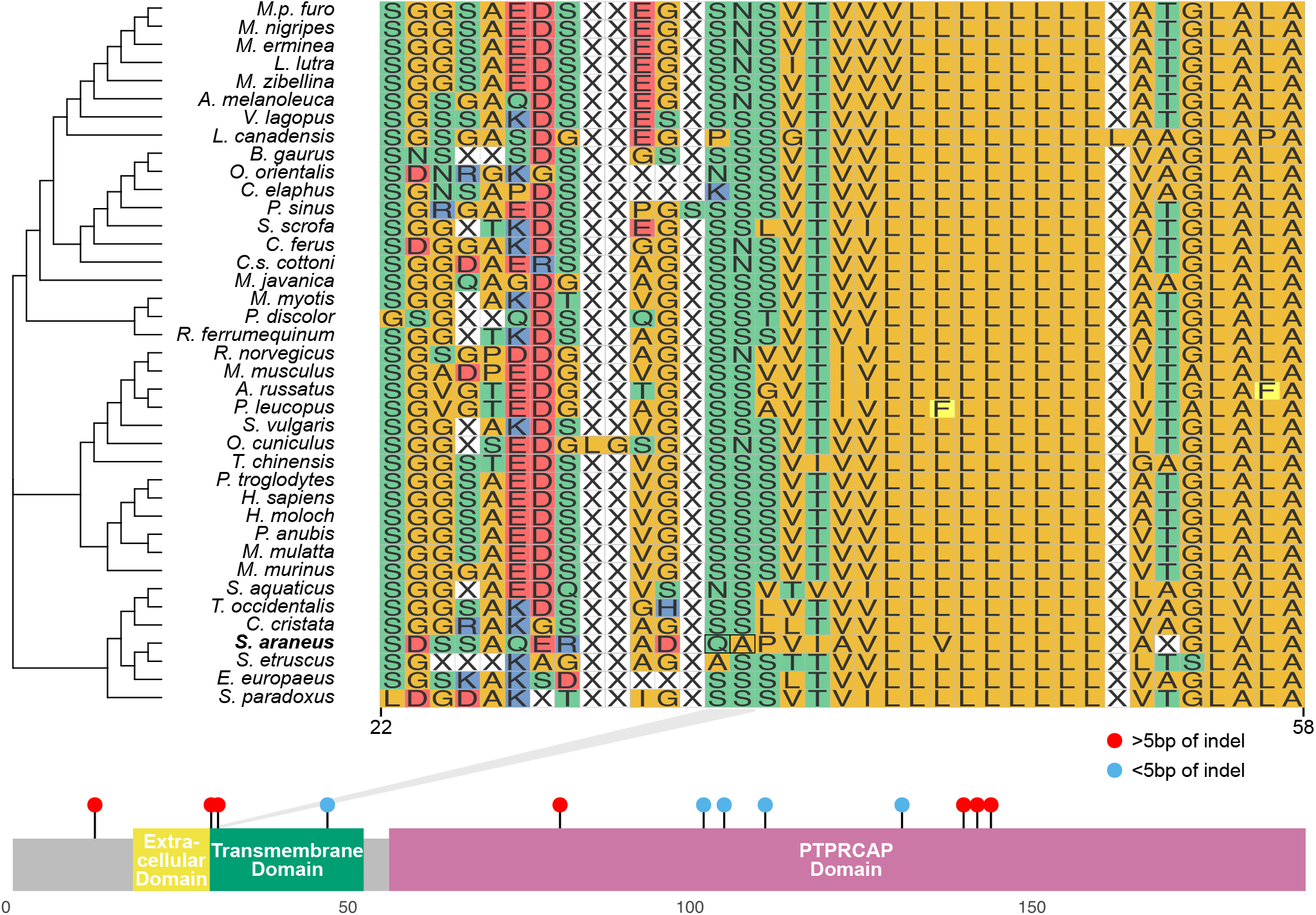
Multiple sequence alignment of the extracellular and transmembrane domains of the positively selected gene, *PTPRCAP* (amino acids 22–58), across 39 mammalian species. Two of the 12 sites identified to be under positive selection by MEME are highlighted, which may have functional consequences for *PTPRCAP* binding capabilities to CD45.

## 4. Discussion

We present the first gene expression characterization of the pancreas of Aselli, an evolutionarily novel haemolymphatic organ in the Eurasian common shew. Our analysis revealed significant differences in gene expression between the pancreas of Aselli and the spleen, with 2,500 upregulated and 3,497 downregulated genes. Pathway enrichment analysis indicated genes upregulated in comparison to the spleen were functionally linked to immunity, with upregulated pathways including primary immunodeficiency, antigen processing and presentation, T cell receptor signaling, natural killer cell-mediated cytotoxicity, and cytokine-cytokine receptor interactions. These pathways are predominantly associated with adaptive instead of innate immunity, suggesting the pancreas of Aselli facilitates antigen-dependent immune responses instead of rapid, non-specific immunity. Alternatively, genes involved in sporphyrin metabolism, or the production and degradation of heme for oxygen transport, were downregulated in the pancreas of Aselli. This evidence aligns well with a previous immunohistological study (*5*) showing that the pancreas of Aselli shares a cellular composition similar to that of a lymph node, containing adaptive immune cells such as T cells, B cell, neutrophils, and macrophages, and plays little role in filtering oxygen-carrying red blood cells, a key function of the spleen.

Functionally, the pancreas of Aselli differs from a typical lymph node in that its medulla is almost entirely composed of plasma cells (*5*), or terminally differentiated B cells reflecting prior immune experience or antigen exposure. This cellular composition is strongly supported by gene expression data, which indicated a regulatory shift toward plasma cell production. We observed minimal changes in the expression of genes associated with B cell maintenance (*PAX5, BCL6, BACH2*) (*35*), with only *BACH2* showing upregulation. These transcriptional repressors play a crucial role in maintaining B cell identity (*35*–*39*) and preventing premature differentiation into plasma cells (*40, 41*) by inhibiting key genes such as *PRDM1* and *XBP1*. In contrast, three key genes involved in plasma cell differentiation – *XBP1, PRDM1*, and *IRF4* – were all significantly upregulated in the pancreas of Aselli. This reinforces this organ’s role as a site for plasma cell rather than naïve B cell storage, as *PRDM1, IRF4*, and *XBP1* drive the transition to antibody-secreting plasma cells (*35*), with *PRDM1* repressing B cell associated genes (*42, 43*), which can be activated by *IRF4* to promote plasma cell fate (*44*). This pattern in the pancreas of Aselli aligns with single-cell sequencing of mouse immune cells, where *PAX5* and *BACH2* are highly expressed in naïve follicular B cells but absent in long lived plasma cells, which instead express *PRDM1, IRF4*, and *XBP1* (*45*). The significant upregulation of plasma cell differentiation genes in the pancreas of Aselli, alongside minimal changes in B cell maintenance genes, highlights its specialized role in long-term plasma cell production instead of the generalized B cell development typical of lymph nodes, and may explain why more plasma cells are found in the lymph nodes of the shrew compared to other species (*5*).

Gene expression evidence suggests the pancreas of Aselli is not a direct analogue of the bursa of Fabricius, supporting histological work (*5*). When comparing the chicken bursa of Fabricius to the spleen, we found that genes associated with naïve B cells (*PAX5, BACH2, BCL6*) were upregulated in the bursa, a pattern that contrasts with the minimal change observed in the pancreas of Aselli (Figure 2). Despite these differences, a comparison of differentially expressed genes across both analyses revealed a greater overlap than expected by chance hypergeometric test, p<0.001) of concordant changes. These similarities can be observed in the plasma cell differentiation genes. Some genes upregulated in the shrew Aselli, such as *IRF4*, were also upregulated in the bursa of Fabricius. These findings indicate that although the pancreas of Aselli and the bursa of Fabricius are not precise functional equivalents, they have evolved independently to enhance adaptive immunity using some similar regulatory mechanisms.

Natural selection may shape some of the key regulatory genes of the pancreas of Aselli compared to other mammals. When comparing genes upregulated in the pancreas of Aselli to those inferred to be under positive selection specific to the shrew (*34*), we did not find more overlap than expected by chance 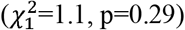. Instead of the continuous evolution, refinement, and innovations of complex immune systems (*46, 47*), or the pancreas of Aselli having increased selection to refine its immune function, as indicated by greater than expected overlap, we find few genomic changes, some perhaps unrelated to this organ’s function. Notably, *PRDM1* was inferred to be under positive selection (Supplemental Figure 1), however, 70% of the sites inferred to be under positive selection were found in the first 100 amino acids, which could represent an alternative start site (*48*). We found *PTPRCAP* was under positive selection (Figure 3), including at the first two amino acids of its transmembrane domain. *PTPRCAP* is typically expressed in lymphocytes (*49*), where it interacts with CD45, a key regulator of antigen receptor–mediated signal transduction and lymphocyte development (*50*). In mice, deletions in the transmembrane, rather than the extracellular domain disrupt *PTPRCAP* binding with *CD45* (*51*), with deficient or knockout mice showing decreased *CD45* expression (*52*) that can, in turn, impair the responses of T and B cell to antigen receptor stimulation (*53*). Yet, *PTPRCAP*-deficient mice have increased cellularity and size of lymph nodes (*52, 54*), suggesting a role in regulating lymphocyte expansion. Evidence of adaptation in the common shrew, particularly within this functional region, may alter antigen presentation while simultaneously supporting lymphocyte development in this novel, enlarged lymph node.

Together, our gene expression analyses support a model in which the pancreas of Aselli and the bursa of Fabricius serve distinct roles in immune development and maintenance, corresponding to the life history of their organisms. The bursa degenerates at sexual maturity (Figure 1B), emphasizing its early-life role in B cell maturation, consistent with the elevated gene expression that sustains mature B cells (*PAX5, BCL6, BACH2*). In contrast, the pancreas of Aselli endures past puberty and throughout the brief lifespan of the shrew. Its gene expression profile, which shows upregulation of plasma cell differentiation genes (*PRDM1, XBP1, IRF4*) in comparison to the spleen, validates its role as a store of plasma cells. These genes likely facilitate the long-term production and secretion of diverse antibodies in response to heavy parasite loads, which continue throughout the shrew’s lifespan, tend to be higher than in other mammals, yet appear to cause no obvious clinical illness (*6, 55*). Both the persistence and gene expression of the pancreas of Aselli suggest the organ functions as a sustained source of adaptive immunity into adulthood.

### 4.1 Limitations

There are methodological limitations in this study. A key challenge in assessing whether the pancreas of Aselli is a direct analogue of the bursa of Fabricius lies in the availability of appropriate comparative sequencing data. To make a robust comparison within *G. gallus*, the bursa and secondary immune organ (the spleen) should ideally be sampled at the same developmental stage with a sufficiently large sample size (at least 4), as low sample sizes increase false positives from high variation in species outside the laboratory (i.e., *PRDM1* in bursa) (*56*). However, after scanning the SRA, the only available high-quality data sets came from two-week-old chicks. This age discrepancy introduces confounding variables when comparing to the pancreas of Aselli, as adaptive immune function can be influenced by developmental stage. Experimental constraints prevent us from fully resolving the mechanisms underlying the formation and evolutionary emergence of this organ. One possibility is that shrews experience a distinct developmental fitness landscape due to their exceptionally short lifespans for mammals (*28*), which may result in novel immune solution. However, our current approach does not allow us to distinguish between genes involved in the organ’s development versus those contributing toward its immune function. Future research integrating embryonic time-series data could provide regulatory insights into the developmental and evolutionary origins of this immune innovation.

## 5. Conclusion

This study provides the first gene expression analysis of the pancreas of Aselli, an evolutionarily novel lymphoid organ, offering new insights into mammalian immune system evolution. Our findings reveal that this organ is functionally distinct from the spleen, with gene expression patterns supporting a specialized role in adaptive immunity, particularly in long-term plasma cell storage. While not a direct analogue of the bursa of Fabricius, as evident by differences in B cell maintenance genes (*PAX5, BACH2, BCL6*) and plasma cell differentiation genes (*IRF4, XBP1, PRDM1*), similarities in regulatory mechanisms suggest independent evolutionary solutions for enhancing adaptive immunity. Additionally, evidence of positive selection in key immune genes, such as *PTPRCAP*, suggests a dynamic interplay between adaptive immunity and natural selection, potentially fine-tuning the function of the pancreas of Aselli to meet the immunological demands of the common shrew. By characterizing the regulatory mechanisms of the pancreas of Aselli alongside selective processes, highlights how evolution and immune function can act synergistically to shape novel adaptations and expand our understanding of mammalian immune system diversity.

## Acknowledgements

We thank Stony Brook University and University of Massachusetts Amherst for key computational resources (SeaWulf and Unity HPCs). BioRender was used to generate Figure 1 and ChatGPT was used to improve manuscript grammar.

## CRediT author contributions

Conceptualization: WRT

Methodology: WRT

Validation: WRT

Formal analysis: WRT

Investigation: WRT

Resources: LMD, TL

Data Curation: WRT, CB

Writing – Original Draft: WRT

Writing – Review and Editing: LMD, CB, DE, TL, YZ, AC, JN

Visualization: WRT

Supervision: LMD

Project administration: LMD

Funding acquisition: LMD, DD, JN, DE, APC

## Data Availability

Supplementary tables, data, results, and code deposited and found on Github https://github.com/wrthomas315/project_aselli. Raw sequencing data located in the NCBI Sequencing Read Archive (BioProject PRJNA941271).

## Figures

**Supplemental Figure 1.**
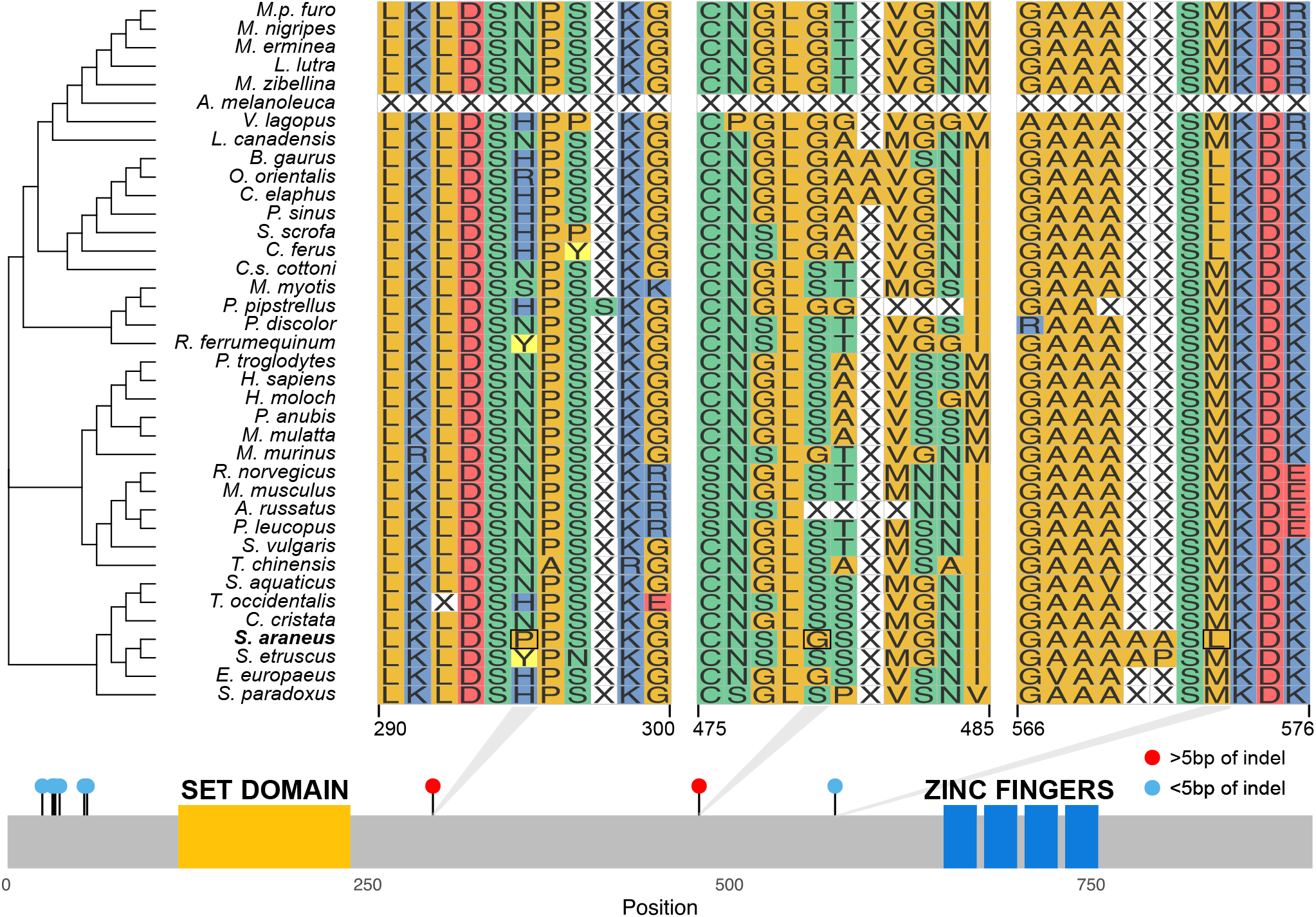
Multiple sequence alignment *PRDM1*. While this gene is indicated to be under positive selection, many (7 of 10) positively selected sites were found within the first 100 amino acids and near indels, reducing confidence in this result.

